# Role of the 5HT_2C_ receptor in anxiety-like behavior in zebrafish (*Danio rerio*)

**DOI:** 10.1101/2025.07.11.664334

**Authors:** Larissa Nunes Oliveira, Loanne Valéria Xavier de Bruce Souza, Brenno Silva Bozi, Bruna Patrícia Dutra Costa, Aurora Rúbria Batista Pantoja, Monica Lima-Maximino, Rachel Coelho Ripardo, Caio Maximino

## Abstract

The present study aims to describe the acute effects of 5-HT_2C_ receptor agonists and antagonists in behavioral tests related to anxiety in adult zebrafish (*Danio rerio*). For this, three groups of fish (*n* = 12/group) were used: a group treated with the drug MK-212 (2 mg/kg), a 5-HT_2C_ receptor agonist; another group treated with the drug RS-102221 (2 mg/kg), a 5-HT_2C_ receptor antagonist; and a third group, the control group, using a vehicle solution. The three groups were exposed to two behavioral tests: novel tank test (NTT) and light-dark preference (LDT). MK-212 produced no effects on the NTT, while RS-102221 decreased geotaxis (d = 0.99, 95%CI[0.3, 1.7]) and erratic swimming (d = 0.87, 95%CI[0.18, 1.56]) in this test; these effects are consistent with a tonic facilitation of defensive behavior in the NTT. Conversely, in the LDT, MK-212 increased scototaxis (d = 0.96, 95%CI[0.25, 1.65]), risk assessment (d = -1.31, 95%CI[-2.02, -0.59]), and thigmotaxis (d = -1.48, 95%CI[-2.21, -0.75]), consistent with an anxiogenic-like effect of this drug. Thus, tonic and phasic effects of the activation of this receptor are observed depending on the type of behavioral test used.

## 1. Introduction

Anxiety is a common and frequent state in human beings, essential for the survival and protection of the individual. However, when it is intense and frequent, it becomes detrimental to the quality of life of this individual, causing discomfort, emotional state, disturbance and distress (Barlow, 2002). Anxiety disorders refer to a group of mental disorders characterized by feelings of anxiety and fear, including generalized anxiety disorder (GAD), panic disorder, phobias, and social anxiety disorder with symptoms that can vary from mild to severe. The duration of symptoms typically experienced by people with anxiety disorders makes it more of an unstable disorder than an episodic one (World Health Organization, 2022).

Seeking to understand the triggering mechanisms of anxiety-like behavior can be a path to a greater understanding of anxiety disorders. Understanding the role of serotonin in the manifestation of anxiety appears to be a promising path to broadening our understanding of it (Maximino, 2012). In the Deakin-Graeff theory, different brain regions have unique and relevant functions for the pathophysiology of anxiety, related to serotonin subpopulations located in the dorsal raphe nucleus and median raphe nucleus (Deakin & Graeff, 1991; Graeff et al., 1997). Thus, serotonin plays a dual role in the regulation of anxiety: anxiogenic substance in the amygdala and anxiolytic in the dorsal periaqueductal gray (Deakin & Graeff, 1991; Graeff et al., 1997) through projections from the raphe to midbrain (e.g., periaqueductal gray area) and forebrain (e.g., amygdala and hippocampus) (Deakin & Graeff, 1991; Graeff et al., 1996). 5-HT, however, is a very complex system, with seven families of receptors found in vertebrates (Bockaert et al., 2010).

One of these families, the 5-HT_2_ family, comprises three subtypes, 2A, 2B, and 2C, that are functionally linked to phospholipases C and A as intracellular transduction mechanisms (Bockaert et al., 2010). Among those subtypes, the 5-HT_2C_ receptor has emerged as a target for the study of anxiety-like behavior. Mora et al. (1997) demonstrated that, in the elevated T-maze, administration of the selective 5-HT_2C_ antagonists SB 200646A and SDZ SER 082 or the mixed 5-HT_2A/2C_ antagonist ritanserin impaired inhibitory avoidance, without affecting one-way escape. This suggests that inhibitory avoidance (a measure of anxiety in this test) is tonically facilitated through the 5-HT_2C_ receptor. In the elevated plus maze, acute administration of the selective 5HT_2C_ agonists MK-212 (de Mello Cruz et al., 2005) or WAY 161503 (Gomes et al., 2010) in Wistar rats selectively reduced exploration of the open arms and increased risk assessment without affecting locomotor activity. WAY 161503 also acts as a discriminative aversive stimulus in taste conditioning and elicits conditioned place avoidance; however, since amphetamine also produced taste aversion to saccharin while simultaneously producing conditioned place preference, the effects of this agonist are likely state-dependent (Mosher et al., 2006). Thus, phasic activation of the 5-HT_2C_ receptor also appears to modulate anxiety-like behavior in rodents. Nonetheless, in zebrafish (*Danio rerio*), MK-212 and WAY 161503 blocked freezing and geotaxis during exposure to a distal threat (conspecific alarm substance, which elicits a fear-like state (Maximino et al., 2019), without effects on post-exposure behavior (potential threat), suggesting that the effects of these drugs are state-dependent (do Carmo Silva et al., 2021). Indeed, MK-212 blocks some of the effects of acute restraint stress on anxiety-like behavior in zebrafish (do Carmo Silva et al., 2021). The selective 5-HT _2C_ antagonist RS-102221, on the other hand, blocked changes in behavior after exposure to CAS (potential threat)(do Carmo Silva et al., 2021), suggesting a tonic facilitation of anxiety-like behavior in this species as well.

These conflicting results suggest that the 5-HT_2C_ receptor has complex effects on threat-motivated behavior across species. While the results from zebrafish seem to point to a phasic modulation of (fear-like) responses to distal threat and a tonic facilitation of (anxiety-like) responses to potential threat, the results from the elevated T-maze seem to indicate that the 5-HT_2C_ receptor does not act on fear-like responses. Nonetheless, both the results from rats and zebrafish seem to position the effects of the 5-HT_2C_ receptor as state-dependent. In the present work, the acute effects of MK-212, a 5-HT_2C_ receptor agonist, and RS-102221, a 5-HT_2C_ receptor antagonist, were evaluated in two commonly used behavioral assays for anxiety-like behavior in zebrafish, the light/dark test and the novel tank test. There is some evidence for different stimulus control in both tests (Kysil et al., 2017; Maximino et al., 2012), with behavior in the first being controlled by an approach-avoidance conflict (and therefore more similar to inhibitory avoidance in the rat elevated T-maze) and behavior in the latter being controlled by escape from the top (and therefore more similar to one-way escape in the rat elevated T-maze).

## 2. Materials and methods

### 2.1 Animals and housing

Adult (>4 month-old) zebrafish (*Danio rerio*) from the longfin phenotype (n = 54) of both sexes, obtained from a colony resulting from crosses between animals purchased from specialized stores. All animals were kept in 50 L tanks (with a density of 80-100 animals per tank) and acclimated in the laboratory for a period of 14 days prior to the experiments, under optimized maintenance conditions. The tanks were filled with dechlorinated water, which was maintained under constant aeration and chemical and mechanical filtration at a temperature of 28 ± 2°C and pH 7.2, according to the standards of care for zebrafish (Lawrence, 2007). The room lighting was provided by fluorescent lamps. The animals were fed twice a day with commercial feed (28% crude protein). Approximately 30% of the water in the tanks was changed daily. The parameters for handling the individuals followed the rulings by the National Council for the Control of Animal Experimentation (Diretriz brasileira para o cuidado e a utilização de animais para fins científicos e didáticos -DBCA. Anexo I. Peixes mantidos em instalações de instituições de ensino ou pesquisa científica, 2017), and the present work was submitted to the Ethics Committee for the Use of Animals (CEUA/Unifesspa) under protocol 23479.020944/2022-04.

### 2.2 Drugs and treatments

Animals were randomly sorted for treatment with vehicle (Cortland’s saline solution (NaCl 124.1 mM, KCl 5.1 mM, Na2HPO4 2.9 Mm, MgSO4 1.9 mM, CaCl2 1.4 mM, NaHCO3 11.9 mM, 1000 units of heparin (Perry et al., 1984)); the 5-HT_2C_ receptor agonist, MK-212 (6-chloro-2-(1-piperazinyl)pyrazine; CAS number #64022-27-1; 2 mg/kg); or the 5-HT_2C_ receptor antagonist RS-102221 (8-[5-(2,4-Dimethoxy-5-(4-trifluoromethylphenylsulfonamido)phenyl-5-oxopentyl]-1,3,8-triazaspiro[4.5]decane-2,4-dione; CAS Number #185376-97-0; 2 mg/kg). These doses were chosen based on their behavioral effects described in adult zebrafish (de Moura et al., 2023; do Carmo Silva et al., 2021). Animals were randomly sorted to drug treatment (*N* = 18/group), using an online randomizer tool (https://www.randomizer.org/). The test subject was initially cryoanesthetized with cold water (∼12 °C); immediately after the appearance of signs of anesthetic plane, it was transferred to a moistened surgical bed where the corresponding drug was injected using a microsyringe (Hamilton 701N, needle size 26, gauge with conical tip 200, 10 μL), according to the protocol described in Kinkel et al. (2010). After injection, the subject was immediately transferred to a recovery tank, consisting of an aquarium measuring 15 cm x 25 cm x 20 cm (width x length x height) with a 10 cm column of dechlorinated water, for 15 minutes. Experimenters and analysts were blinded to drug treatment by using coded vials; the blind was removed only after statistical analysis. After the 15 minutes of recovery, animals were randomly sorted across test orders, with half of the animals being tested first in the novel tank test (NTT) and then in the light/dark test (LDT), both described below, and the other half being tested first in the LDT and then in the NTT. Each fish performed both tests sequentially in order not to exceed the time and the drug to be eliminated before the end of testing (Figure 1).

**Figure 1.**
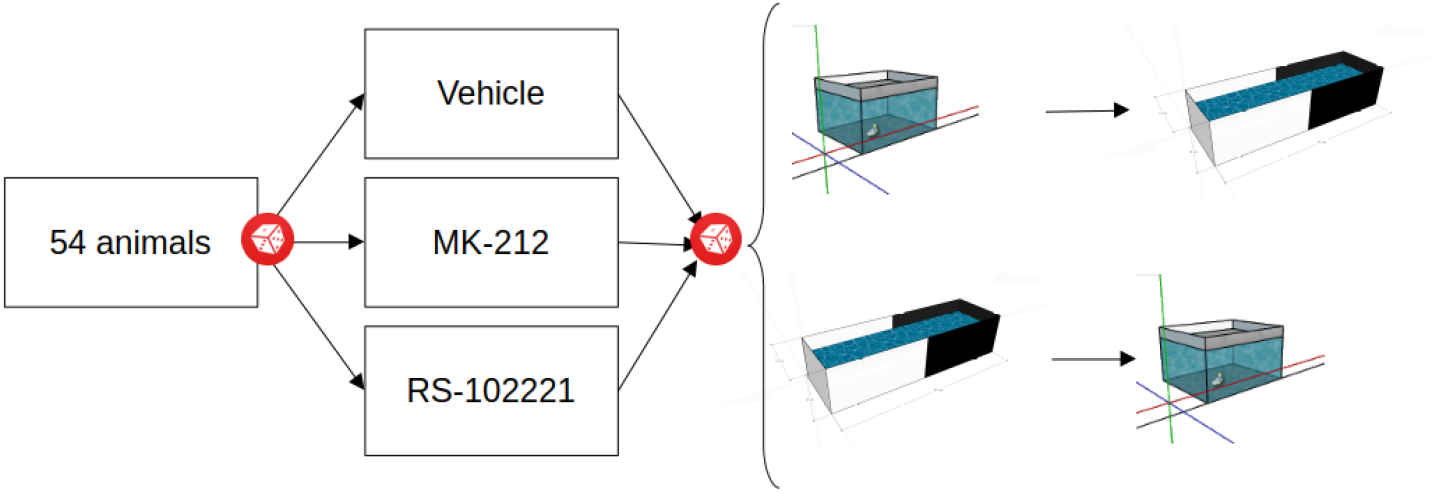
Experimental design. Animals were randomly assigned (dice icon) to drug treatment (vehicle, MK-212, or RS-102221), and later randomly assigned to test order. More details can be found in the Methods section.

### 2.3 Experimental conditions

The tasks described below aim to evaluate parameters related to locomotion/exploration and anxiety, and were carried out between 10:00 and 15:00, in order to respect the photoperiod of the individuals. The lighting of the room is a critical variable, as it can alter the behavior of the fish (Facciol et al., 2017, 2019), so immediately before each test, its measurement was measured using a lux meter, resulting in an average of 518.7 lux ± 39.7 above the tank. In order to block possible noise interference, the experiment was performed under constant Gaussian white noise, producing an average of 63.2 dB ± 3.96 above the tank.

#### 2.3.1 Experiment 1: Novel Tank Test

This task aims to evaluate zebrafish vertical locomotion/exploration, as well as fear/anxiety behaviors related to responses to novelty (Bencan et al., 2009; Cachat et al., 2011; Egan et al., 2009; Levin et al., 2007). In this first task, the animal freely explored a novel tank for 6 minutes, and the novelty of the environment elicited escape responses from the top (Maximino et al., 2012). The test apparatus consisted of an aquarium measuring 15 × 25 × 20 (width x length x height) filled with water up to its tip (Maximino et al., 2013). All activity was recorded on film. The video analysis was performed using TheRealFishTracker v.0.4.0 software (https://www.dgp.toronto.edu/~mccrae/projects/FishTracker/). In the analysis, the parameters considered were based on Kalueff et al. (2013) with the corresponding definition provided:

1-Geotaxis (duration): Time spent in the lower third of the aquarium (ZBC1.46);

2-Surfacing (duration): Time spent in the upper third of the tank;

3-Erratic swimming (absolute turning angle): zigzag movements characterized by rapid changes in linear and angular acceleration in short sequences, measured by the absolute turning angle (ZBC1.51);

4-Swimming speed (cm/s): the speed was evaluated in order to investigate whether the drug had a sedative effect on the animal;

#### 2.3.2 Experiment 2: Light/Dark Test

The scototaxis behavior in zebrafish, characterized by the preference for dark environments over light environments, has been described and validated as a model for the evaluation of anxiety-like behavior for the species (Araujo et al., 2012; Maximino et al., 2010, 2011, 2014). The subject was transferred into the central compartment of the light/dark test apparatus for 3 minutes, following the protocol established by the research group (https://www.protocols.io/view/light-dark-preference-test-for-adult-zebrafish-dan-bp2l65yzgqe5/v2). The apparatus consisted of an acrylic tank (15 cm × 10 cm × 45 cm height × width × length) that is divided equally into black and white halves. The walls and bottom were either black or white to ensure uniform substrates for each compartment. The water column height was maintained at 10 cm, resulting in a final volume of 4.5 liters. The water used was dechlorinated. The colored material chosen should not be reflective to avoid the tendency to shoal and behave in relation to their own reflection. The tank contains central sliding doors, colored with the same color as the side of the tank, thus defining a colorless central compartment. After 3 minutes for the subject to re-acclimate, the central sliding doors were removed and the test and its image recording are started. This test lasts 15 minutes. The collected data were analyzed using the X-plo-rat software (https://github.com/lanec-unifesspa/x-plo-rat; Tejada et al., 2018).

In the analysis, the parameters considered were based on Kalueff et al. (2013) with the corresponding legend provided:

1. Scototaxis (duration): time in which the animal remains in the white compartment (ZBC1.137);
2. Number of entries (frequency): moment in which the animal enters the white compartment for more than 5 seconds (ZBC1.54);
3. Erratic swimming (frequency): zigzag movements characterized by rapid changes in linear and angular acceleration in short sequences, measured by the absolute turning angle (ZBC1.51);
4. Risk assessment (frequency): total or partial entry of the animal into the white compartment lasting less than 5 seconds (n/a);
5. Thigmotaxis (percent): swimming only along the edges of the test apparatus (ZBC1.173)

### 2.4 Statistical Analysis

The results obtained by the behavioral tests were analyzed through one-way analysis of variance (ANOVA), followed by Tukey’s post hoc test. Effect sizes were calculated as ω^2^. The data was represented as Gardner-Altman plots (Ho et al., 2019), with individual datapoints plotted on the upper axes. On the lower axes, each effect size (Cohen’s d; Cohen, 1988) was plotted as a bootstrap sampling distribution. Bootstrap resamples were taken using Efron’s (1987) method; the confidence interval was bias-corrected and accelerated. Plots were made using R 4.1.2, using the dabestr package (v. 2023.9.12; https://github.com/ACCLAB/dabestr). Post-hoc power calculations were made using the pwr package (v. 1.3-0; https://cran.r-project.org/web/packages/pwr/index.html) in R, and the power of each analysis is reported.

## 3. Results

### 3.1 Novel Tank Test

In the novel tank test, a significant effect of drug was found for geotaxis (F[_2,51]_ = 6.02, p = 0.004, ω^2^ = 0.16; Figure 2A). Post-hoc tests found that MK-212 had no effect in this variable (p = 0.99; d = -0.01, 95%CI[-0.67, 0.67]), while RS-102221 decreased geotaxis (p = 0.011; d = 0.99, 95%CI[0.3, 1.7]). No effects were observed on surfacing (F_[2, 51]_ = 2.13, p = 0.13, ω^2^ = 0.04; Figure 2B). Significant effects of drug were found for erratic swimming (F_[2, 51]_ = 3.74, p = 0.031, ω^2^ = 0.09; Figure 2C); post-hoc tests found that RS-102221 decreased erratic swimming (p = 0.031, d = 0.87, 95%CI[0.18, 1.56]), while MK-212 did not produce an effect (p = 0.12, d = 0.67, 95%CI[-0.01, 1.35]). No effects were observed on swimming speed (F_[2, 51]_ = 1.85, p = 0.168, ω^2^ = 0.03; Figure 2D).

**Figure 2.**
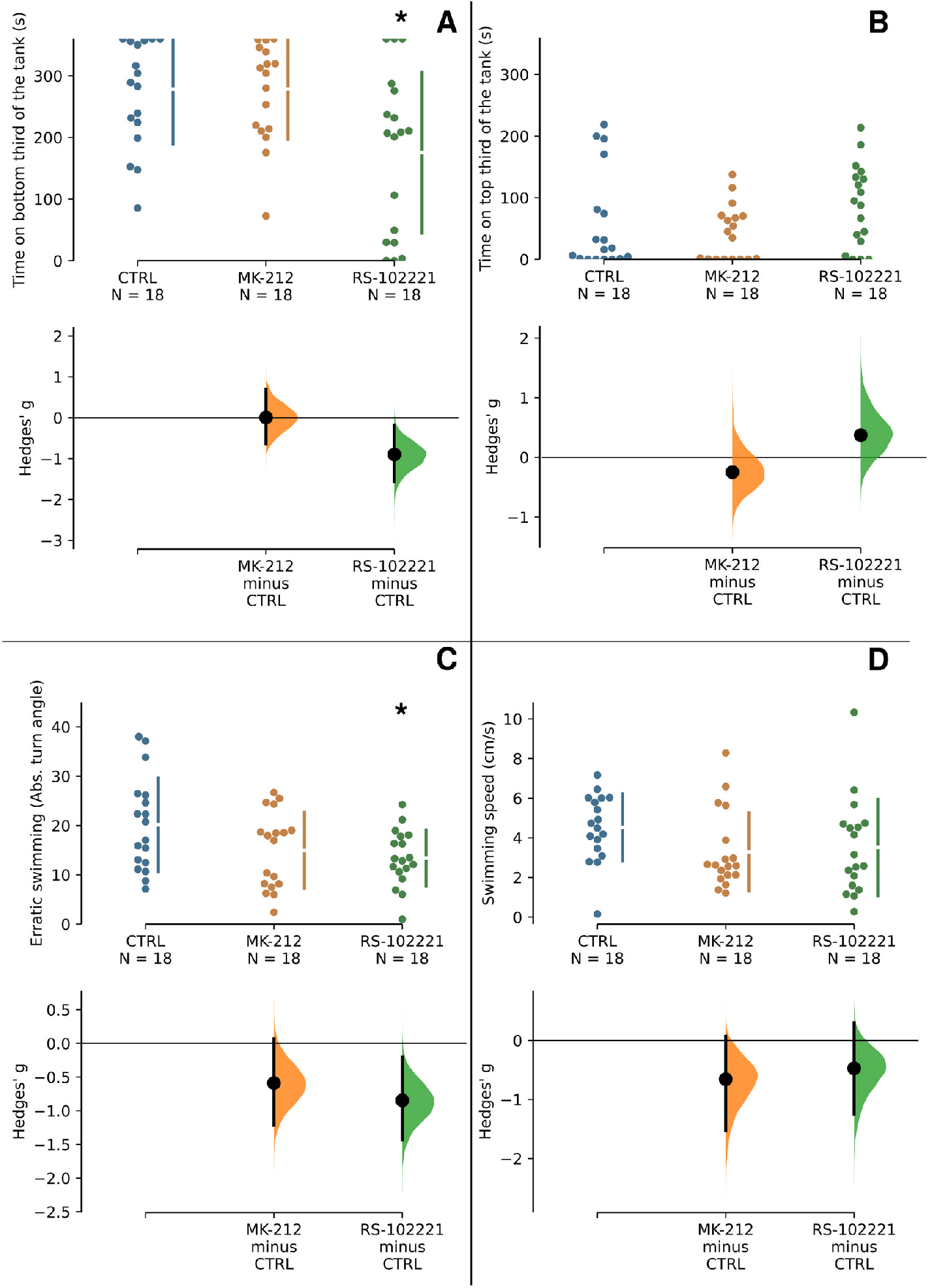
Treatment with RS-102221, but not MK-212, produce anxiolytic-like effects in the novel tank test (NTT). Effects of treatment with vehicle, MK-212 (2 mg/kg)), or RS-102221 (2 mg/kg) on the (A) Geotaxis, (B) Surfacing, (C) Erratic swimming, and (D) Swimming speed during the Novel Tank Test behavioral test. The mean difference between the MK-212 and RS-102221 groups with shared control (CTRL) is shown in the Cumming estimate plot. Raw data are plotted on the upper axes. On the lower axes, effect sizes (Hedges’ *g*) are plotted as bootstrap sampling distributions. Each effect size estimate is represented as a point. Each 95% confidence interval is indicated by the tails of the vertical error bars. Asterisks (*) mark statistically significant differences against CTRL.

### 3.2 Light and Dark Test

A main effect of drug was found for time on white (F_[2, 51]_ = 5.68, p = 0.006, ω^2^ = 0.15; Figure 3A). Post-hoc tests found that MK-212 significantly decreased time on white (p = 0.016; d = 0.96, 95%CI[0.25, 1.65]), while RS-102221 had no effect in this variable (p = 0.0994; d = -0.03, 95%CI[-0.7, 0.63]). No effects were observed for entries on white (F_[2, 51]_ = 1.94, p = 0.154, ω^2^ = 0.03; Figure 3B). A significant effect of drug was found for risk assessment (F_[2, 51]_ = 12.11, p < 0.001, ω^2^ = 0.29; Figure 3C); post-hoc tests found that MK-212 significantly increased risk assessment (p < 0.001; d = -1.31, 95%CI[-2.02, -0.59]), while RS-102221 had no effect (p = 0.813; d = 0.2, 95%CI[-0.47, 0.87]). No effects were found on erratic swimming (F_[2, 51]_ = 2.28, p = 0.113, ω^2^ = 0.05; Figure 3D). A significant effect of drug was found for thigmotaxis (F[2, 51] = 14.27, p < 0.001, ω^2^ = 0.33; Figure 3E). Post-hoc tests found that MK-212 increased thigmotaxis (p <0.001; d = -1.48, 95%CI[-2.21, -0.75]), while RS-102221 had no effect (p = 0.989; d = 0.11, 95%CI[-0.56, 0.78]).

**Figure 3.**
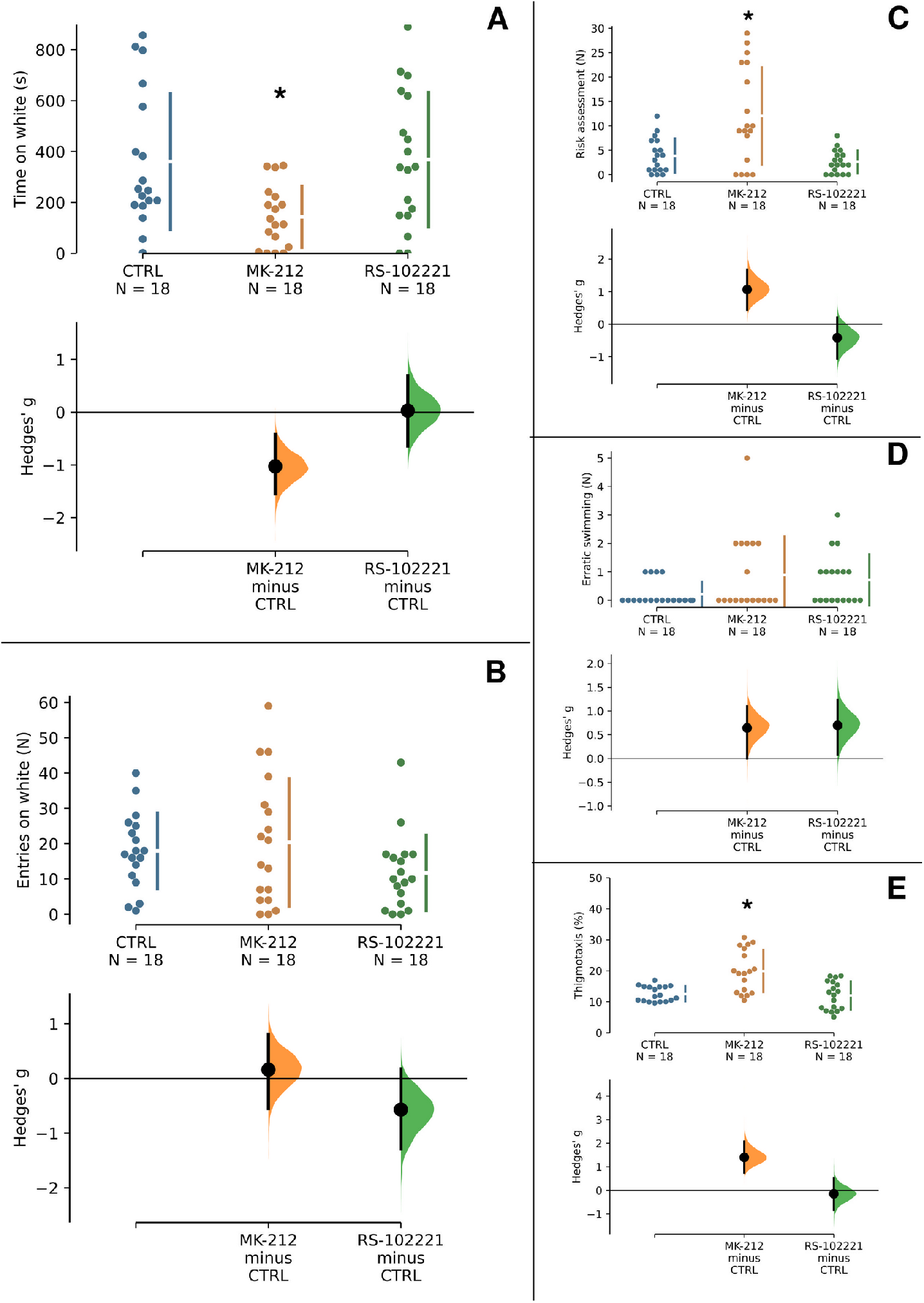
Treatment with MK-212, but not RS-102221, produces anxiogenic-like effects in the light/dark test (LDT). Effects of treatment with vehicle, MK-212 (2 mg/kg), or RS-102221 (2 mg/kg) on the (A) Scototaxis (time on white), (B) Entries on the white compartment, (C) Risk assessment, (D) Erratic swimming, and (E) Thigmotaxis. The mean difference between the MK-212 and RS-102221 groups with a shared control (CTRL) is shown in the Cumming estimate plot above. Raw data are plotted on the upper axes. On the lower axes, effect sizes (Hedges’ *g*) are plotted as bootstrap sampling distributions. Each effect size estimate is represented as a point. Each 95% confidence interval is indicated by the tails of the vertical error bars. Asterisks (*) mark statistically significant differences against CTRL.

## 4. Discussion

The present work investigated the acute effects of the 5-HT_2C_ receptor ligands MK-212 (agonist, 2 mg/kg) and RS-102221 (antagonist, 2 mg/kg) on parameters of anxiety-like behavior in adult zebrafish in the novel tank test (NTT) and the light/dark test (LDT). In general, RS-102221 decreased geotaxis and erratic swimming in the NTT, while MK-212 increased scototaxis, risk assessment, and thigmotaxis in the LDT. RS-102221 did not affect behavior in the LDT, and MK-212 did not affect behavior in the NTT.

Geotaxis is the main index of defensive behavior in the NTT; in general, anxiogenic and/or stress-inducing drugs and treatments increase the time spent at the bottom of the tank, with a concomitant increase in erratic swimming and freezing, while anxiolytic drugs and treatments produce the opposite pattern (Cachat et al., 2011; Egan et al., 2009). In this test, RS-102221 decreased geotaxis and erratic swimming, without effects on surfacing, freezing, or swimming speed. These effects are consistent with a tonic, but not phasic, facilitation of defensive behavior in the NTT, and may be an indication of an anxiolytic effect of the antagonist. This effect is in agreement with data found in Do Carmo Silva et al. (2021) where the same dose blocked increases in geotaxis and erratic swimming after exposure to the alarm substance, indicating this possible anxiolytic effect of the drug. In another study with the NTT, ketanserin, a non-selective 5-HT_2_ receptor antagonist, increased geotaxis and erratic swimming, suggesting an anxiogenic-like effect (Nowicki et al., 2014); while ketanserin has affinities at the nanomolar range for all 5-HT_2_ receptors, at least in human receptors its affinity is higher for the 5-HT_2A_ than the other receptors in this family (https://www.guidetopharmacology.org/GRAC/LigandDisplayForward?tab=biology&ligandId=88), suggesting that the anxiogenic-like effect of this drug is mediated by the first receptor. Moreover, ketanserin also binds alpha-adrenergic receptors at the nanomolar range, which further complicates the comparison of the present results with those reported by Nowicki et al., (2014). Nonetheless, MK-212 did not produce an effect in the present experiments, which is in disagreement with rodent data on both the elevated T-maze (in which antagonists impair inhibitory avoidance, but not one-way escape) and in the elevated plus-maze (in which agonists produce an anxiogenic-like effect)(de Mello Cruz et al., 2005; Gomes et al., 2010; Mora et al., 1997).

In contrast, during exposure to the LDT, the antagonist did not produce a behavioral effect, while MK-212 increased scotoxaxis, risk assessment, and thigmotaxis. Scototaxis in adult zebrafish, characterized by the preference for dark environments over light environments, has been described and validated as a model for the evaluation of anxiety-like behavior for the species (Araujo et al., 2012; Maximino et al., 2010). Risk assessment and thigmotaxis are also sensitive to both anxiolytic-like and anxiogenic-like drugs and stressful stimuli (Lima et al., 2016; Maximino et al., 2014; Quadros et al., 2016). The effects of MK-212 are consistent with a phasic, but not tonic, facilitation of defensive behavior in the LDT by the 5-HT_2C_ receptor, and may be an indication of an anxiogenic effect of the agonist.

Taken together, results from both the NTT and LDT show concordant effects of agonists and antagonists in behavior in these assays, although the pattern of modulation of receptor activation is different in each test (phasic vs. tonic). An effect of the agonist suggests phasic modulation, since it is independent of 5-HT levels; an effect of the antagonist in the absence of an agonist suggests tonic modulation, since, for an antagonist to produce a behavioral effect, it must displace 5-HT that is already binding to receptors, and therefore a serotonergic tonus must be acting. Considering that both tests are usually considered parameters for evaluating anxiety-like behavior, this result could indicate a difference in the sensitivity of the tests for evaluating drug effects; however, the meta-analysis by Kysil et al. (2017) shows that both tests have similar sensitivity to drugs and toxicants in zebrafish, emphasizing their usefulness in development for neurobehavioral research that studies them. Stimulus control in both tests might explain the different effects of MK-212 and RS-102221 in the present experiments; in the NTT, geotaxis is controlled mainly by escape of the top region of the tank, while in the LDT scototaxis is controlled mainly by an approach-avoidance conflict (Maximino et al., 2012). This is also consistent with results from experiments with distal threats (conspecific alarm substance), in which 5-HT_2C_ receptor agonists blocked freezing and geotaxis during exposure (do Carmo Silva et al., 2021). This suggests that phasic activation of the 5-HT_2C_ receptor is panicolytic and anxiogenic in zebrafish.

Another important difference in the tests is related to brain 5-HT release dynamics. When compared to handling, exposure to the LDT increases 5-HT levels in the extracellular fluid of the zebrafish brain, while exposure to the NTT does not produce this effect; when tissue 5-HT levels are assessed, exposure to the NTT increases 5-HT in the midbrain, while exposure to the LDT increases 5-HT in the hindbrain and forebrain (Maximino et al., 2013). These effects are not related to glucocorticoid release dynamics, since exposure to the NTT or the LDT both increase whole-body cortisol (Kysil et al., 2017). Zebrafish possess two copies of the gene coding for the 5-HT_2C_ receptor, *htr2cl1* and *htr2cl2* (Sourbron et al., 2016); the binding site is conserved in both isoforms (de Moura et al., 2023), and therefore it is not possible to know which isoform is being activated or blocked by MK-212 or RS-102221. While the localization of 5-HT_2C_ receptors in the adult zebrafish brain is unknown, *htr2cl1* is expressed in the larval olfactory bulb, dorsal thalamus, eminentia thalami, pretectum, posterior tuberculum, hypothalamic area, and medulla oblongata (Schneider et al., 2012). Differences in circulating levels of 5-HT elicited by each test, a function of the differences in stimulus control, could be responsible for the different effects of 5-HT_2C_ agonists and antagonists in the NTT and LDT.

Although both tests indicate behavioral effects of 5-HT_2C_ receptor agonists and antagonists, the tests differ in stimulus control. Therefore, further research is needed involving the 5HT_2C_ receptor and anxiety-like behavior with its excitation phases, considering both tests.

## Acknowledgments

This work was supported by a Conselho Nacional de Desenvolvimento Científico e Tecnológico (CNPq/Brazil) productivity grant to CM (#302998/2019-5). LNO an LVXBS were supported by Fundação Coordenação de Aperfeiçoamento de Pessoal de Nível Superior (CAPES/Brazil) scholarships.

